# Months-long stability of the head-direction system

**DOI:** 10.1101/2024.06.13.598909

**Authors:** Sofia Skromne Carrasco, Guillaume Viejo, Adrien Peyrache

**Affiliations:** Montreal Neurological Institute, McGill University, Montreal, QC, Canada; Center for Computational Neuroscience, Flatiron Institute, New York, NY, USA

## Abstract

Spatial orientation is a universal ability that allows animals to navigate their environment. In mammals, the head-direction (HD) system is an essential component of the brain’s navigation system, yet the stability of its underlying neuronal code remains unclear. Here, by longitudinally tracking the activity of the same HD cells in freely moving mice, we show that the internal organization of population activity in the HD system was preserved for several months. Furthermore, the HD system developed a unique mapping between its internal organization and spatial orientation in each environment. This was not affected by visits to other environments and was stabilized with experience. These findings demonstrate that stable neuronal code supports the sense of direction and forms long-lasting orientation memories.

## INTRODUCTION

Although animals are capable of life-long memories, brain representations change over time (*1*–*6*). This “representational drift” has been observed in sensory systems, where the tuning of individual neurons to e.g. olfactory (*2*) and visual inputs (*3–5*) changes over days. This is also the case in higher-order systems such as the hippocampus, where the spatial tuning of place cells (*7*) changes across days (*1*, *8–11*), and even between successive trials (*12*). In contrast to this temporal instability of single neurons, distributed representations pertaining to environment identity (*13*) or precise behavioral correlates (*14*) can be stably decoded over time.

These observations beg the question of the stability of the neuronal population code upstream the hippocampus. To address this question, we study the post-subiculum (PoSub), the primary cortical stage of the HD system where most neurons encode the animal’s head-direction (*15*, *16*). The HD system is believed to be governed by attractor dynamics (*17–21*), which constrain neuronal population activity to lie on a one-dimensional ring. In mammals, this was suggested by the stability of internal population dynamic across brain states (*16*, *18*, *22*) yet these observations have only been made within single days of recordings. Furthermore, HD neurons map their internal states to the animal’s direction in an allocentric reference frame by integrating external sensory inputs (*23*, *24*). This mapping is rapidly learned during the first exposure to an environment (*25*). However, the stability of these orientation memories across visits is unclear (*15*). We thus tested the hypothesis that the population of HD neurons maintains their coordination and their alignment to the external world for extended periods, across days.

## RESULTS

To determine the long-term stability of the HD signal, we implanted 5 mice with a 1.0 mm GRIN lens over the PoSub (Fig. 1A, Methods). We detected registered regions of interest (ROIs) in each recording (21 – 286 ROI, mean 95 ±56 s.t.d.) in a total of 388 recording sessions.

**Figure 1:**
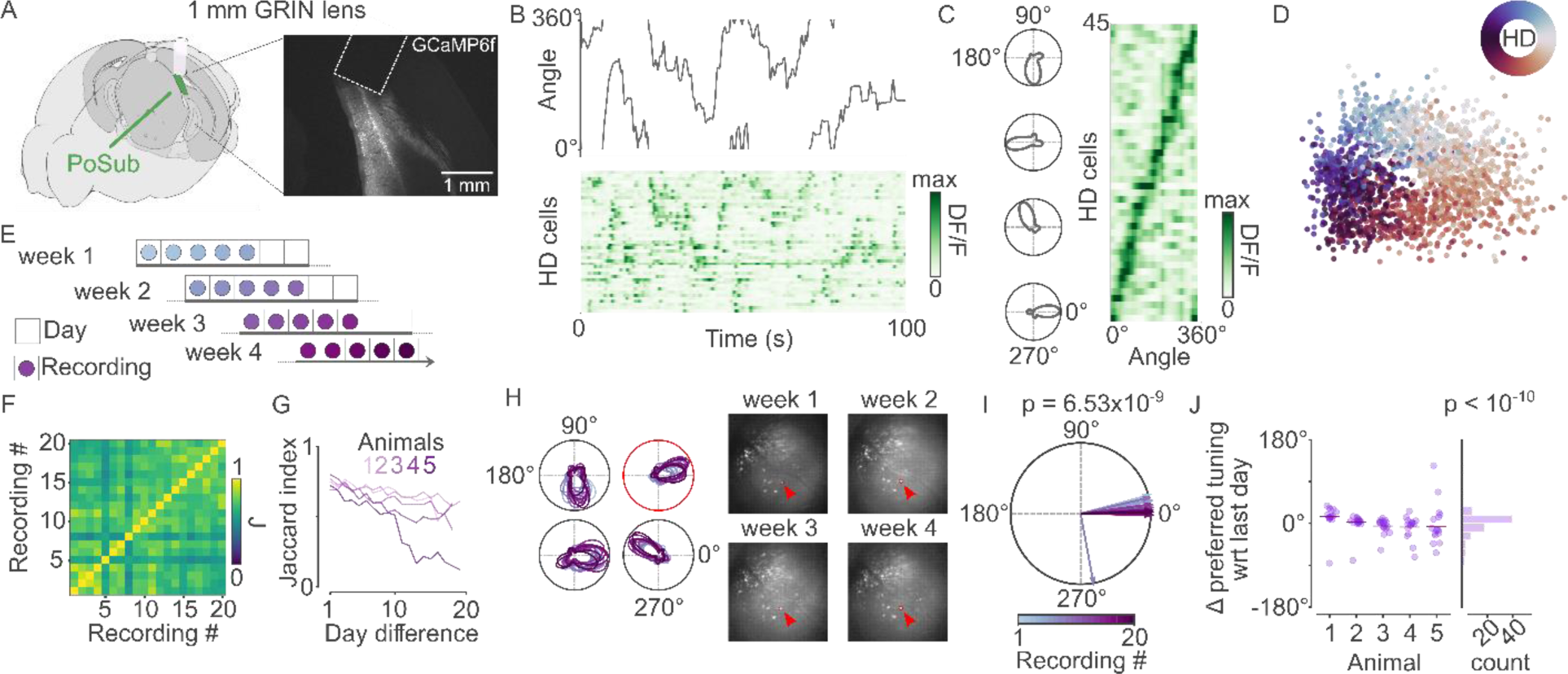
Long-term stability of PoSub HD cell tuning in the same environment. A) Diagram of GRIN lens position above the PoSub (left) and example histology (right) illustrated with brainrender (*27*). B) Example recording of animal head-direction during freely moving behavior in a circular environment (top) and HD cell population fluorescence (bottom). C) Example tuning curves of 4 HD cells in polar coordinates (left) and color-coded tuning curves of all HD cells simultaneously detected (right, same data as B). D) Low-dimensional projection of HD cell population calcium transients from an example recording session (see Methods), revealing the ring-shape topology of the HD cell ensemble. E) Recording protocol of the four week-long tracking (20 sessions total) of ROIs in a circular environment. F) Day-to-day Jaccard index for an example animal, showing the similarity of HD cell registration across sessions. G) Average Jaccard index as a function of duration between pairs of sessions for all animals. H) Tuning curves of four example ROIs from the same animal across all recording sessions where the ROIs were detected (left) and spatial footprint of the circled ROI in four sessions across the four weeks of the recording protocol. I) Population average of across-session angular difference in preferred direction, relative to the last session for one example animal (V-test, not different from 0°, p=7×10^-9^, n=18 sessions). J) Distribution of population-wise angular difference relative to the last session for all five mice (V-test, not different from 0°, p<10^-10^, n=84 sessions).

We first characterized the tuning of single putative units over multiple visits in the same environment. ROIs in the PoSub showed strong tuning to the animal’s head-direction (Fig. 1B, C). Low-dimensional projection of ROI population activity using Isomap (see Methods) revealed a ring topology (Fig. 1D), as expected for a population of HD cells (*18*). We then registered ROIs across the whole recording protocol (20 visits over four weeks, Fig. 1E) (*26*) and identified HD cells as ROIs showing strong tuning to head direction and being registered across multiple days (see Methods). The session-to-session registration quality was evaluated as the overlap in HD cell detection using the Jaccard index (see Methods) (Fig. 1F,G). In an example animal, the tuning of single HD cells was highly preserved across days (Fig. 1H, I), and the same observation was made across all animals (Fig. 1J). Finally, angular offset between HD cell pairs was also highly preserved across days (Supplementary Fig. 2E). However, it remained unclear whether this was a property of the system or a by-product of the HD tuning stability of individual HD cells.

We then asked whether HD cells preserve their pairwise angular difference across days and environments. To this end, we recorded animals in four different environments for two weeks in each environment (8 weeks total) (Fig. 2A). ROIs were detected and monitored throughout the recording protocol (Fig. 2B). In an example animal, HD cells showed environment-specific alignment and, importantly, preserved their pairwise angular difference (Fig. 2C,D). We observed the same tendency of preserved pairwise angular difference in preferred directions across all sessions and animals (Fig. 2E, Supplementary Fig. 4B,C; p<10^-10^, K-S test, n=214,126 pairs). We confirmed these imaging results with electrophysiological recordings (Supplementary Fig. 5). Furthermore, the absolute alignment of the HD cell population within each environment was preserved for the two weeks of visits (Supplementary Fig. 3C), as initially observed for four weeks in the circular environment (Fig. 1).

**Figure 2:**
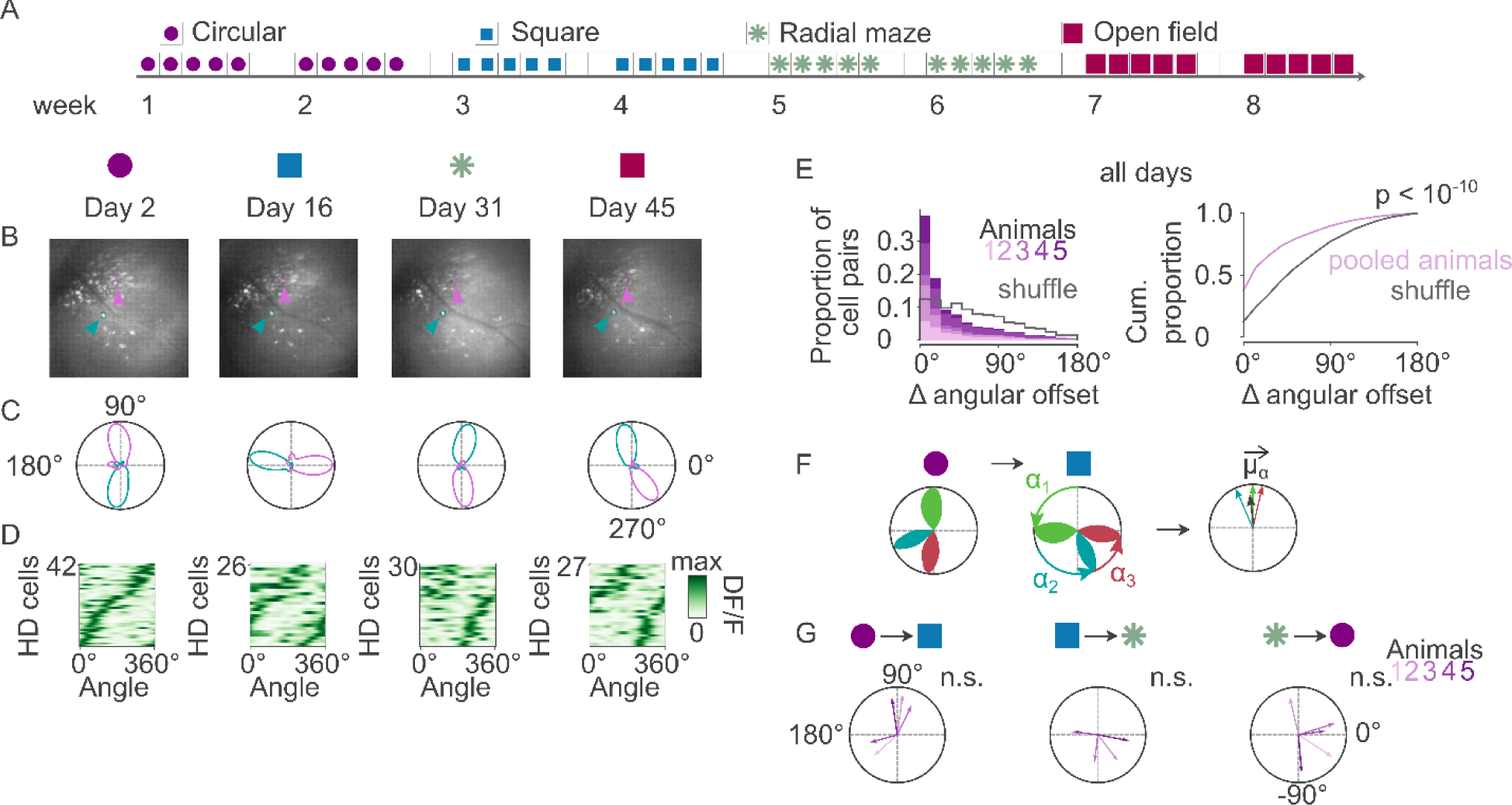
Long-term stability of HD cell population organization across different environments. A) Recording protocol of the eight-week-long tracking (40 sessions total) of ROIs across four different environments. B) Example field of views for recordings across all four environments with two identifiable HD cells. C) Example tuning curves in polar coordinates of the two HD cells in B. D) Tuning curves of all HD cells detected in the example days in B and C, sorted by Day 2. E) Difference in angular offset across all HD cell pairs and all recording pairs in the recording protocol, shown for individual animals (left) and for the pooled population (right) relative to control datasets of shuffled cell pair identities (p < 10^-10^, Kolmogorov-Smirnov two-sample test, n= 214,126 pairs). F) Evaluation of remapping between environments. For each HD cell, we first computed the average angular difference in the preferred direction of each cell across two environments (left). The alignment of the resultant vector of all HD cell angular offsets between environments defined for each animal the angular remapping between environments (µ_α_), and its mean vector length defined the coherency of this remapping. G) Polar representation of the average remapping between pairs of environments for each animal relative to control (shuffled cell identities). The three environments were located in the same room, and alignment was measured with respect to the absolute coordinates of the room. (p > 0.05 for each pair of environments; Rayleigh’s test; n = 5 animals).

The four week-long recording in the circular environment suggested that the absolute alignment of the HD system is maintained for extended periods (Fig. 1). However, it is unclear whether the HD system was explicitly aligned to each environment or with respect to global coordinates. The first three environments of the protocol were in the same experimental room, and the HD system alignment was measured with respect to the absolute coordinates of the room. We thus computed the average remapping of the HD cell population across environments for each animal (Fig. 2F). Interestingly, this remapping was random across animals. While the internal structure of the HD system was maintained between environments, its absolute alignment was specific to each environment. Overall, these observations suggest that HD cells are part of a rigidly configured system that rapidly acquires a new and random alignment when visiting an environment for the first time and maintain this alignment for subsequent visits.

The acquisition of a new absolute alignment in each environment and maintenance of that alignment during two weeks of recordings suggests that the HD system maintains a memory of its mapping to the external world in each environment (Supplementary Fig. 3C). It further begs the question of the long-term stability of this memory. To address this question, we added a week of probe trials after eight weeks of visits (Fig. 2), during which each environment was visited once again (one environment per day) (Fig. 3A). An example HD cell, as well as nearly all the other HD cells recorded simultaneously, showed similar tuning curves in the probe trials and the last visit of the same environment during the 2-week long exploration (Fig. 3B,C). Because environments were initially visited one after the other (Fig. 2), each environment was associated with a different delay between the probe trial and the last visit. For each delay (i.e. each environment) and each animal, we computed the angular remapping of the HD cell population (as in Fig. 2G). The remapping was not different from 0° for delays from 5 days to 4 weeks, although a drift was observed at a six-week interval (Fig. 3D,E). These findings suggest that, in addition to showing stable internal dynamics over a long period, the HD system forms long-term memories of orientation to the external world, which are stable for up to four weeks.

**Figure 3:**
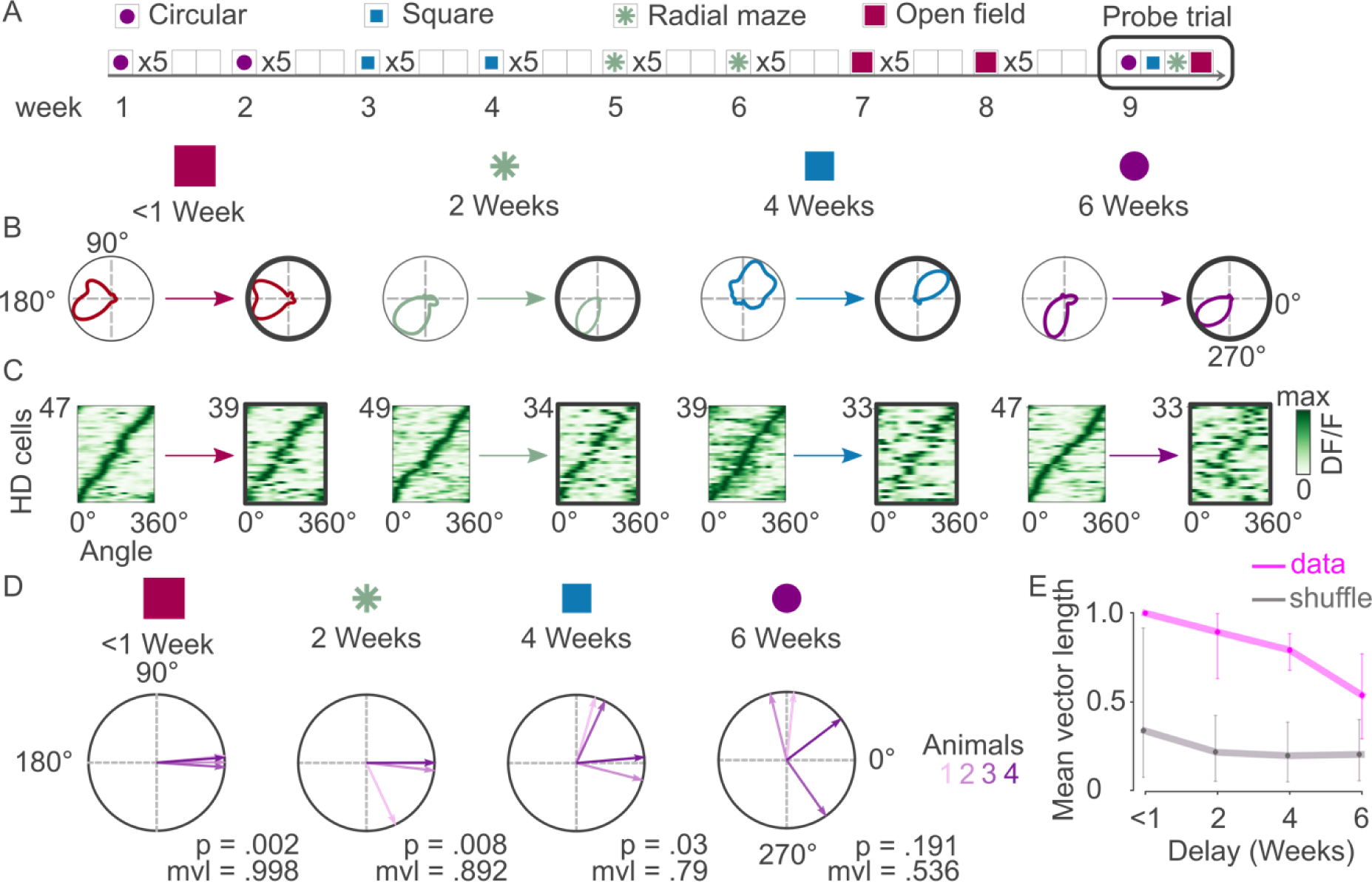
Long-term memory of orientation. A) Memory of orientation was tested in a one-week probe trial following the recording protocol in the four different environments (Fig. 2). B) Tuning curves of an example HD cell recorded during the last visit and probe trials in the four different environments. C) Tuning curves of the HD cell population were recorded on the same days as the HD cell in B and sorted by the initial exploration. D) Change in the preferred direction of the HD cell population in the probe trial with respect to the last visit of that environment during the two-week exploration protocol for each animal. Remapping was not different from 0° across animals for delays from 5 days to 4 weeks, but significantly different from zero after a delay of 6 weeks (V-test, n=4 animals). E) Mean vector length of animal remapping angles (i.e. probe versus last visit of each environment) as a function of delay between visits compared to controls (shuffled HD cell-identity). Error bars in controls depict 95^th^ percentile of the shuffle distribution while error bars in data depict the 95% confidence interval.

While HD cells seem to form a rigidly organized population that is stably aligned over successive visits to the same environment (Fig 1), the slow drift in orientation memory observed with increasing delays between consecutive visits (Fig. 2D,E) could result from two different processes: either an interference with experiences in other environments (hypothesis #1) or a drift in orientation memory accumulating over time between consecutive visits of an environment (hypothesis #2). We tested the first hypothesis by adding three weeks of probe trials to the one following the 8-week long recording protocol of the four different environments (Fig. 3A). The order of the environments was different in each week of probe trials. We observed a strong stability of orientation memory across animals despite interleaving experiences in other environments (Fig. 3B, C). This observation rules out that experiences in other environments interfere with an existing orientation memory.

We then tested whether the delay between two consecutive visits to a given environment could explain the drift in orientation memory. To this end, animals visited two environments on a given day (8 hours apart). It should be noted that animals were already highly familiar with both environments during these visits. Then, animals visited one of the environments every week while they reexperienced the other one only four weeks later. Unlike in the environment visited weekly, we observed a significant drift in orientation memory across animals for the environment that was only visited four weeks apart. Hence, frequent experience in an environment stabilizes orientation memory.

## DISCUSSION

We found that the organization of population activity in the PoSub HD system is stable over time and across environments. Furthermore, the absolute alignment of the HD system in a specific environment was retained when the animal was re-exposed to that environment every week and slowly drifted otherwise.

Our findings that the organization of the HD cell population is rigid across months extend earlier reports that HD cell pairs shift their preferred direction coherently (*23*, *25*) and preserve their mutual coordination across all brain states, including in sleep (*18*, *22*). As the HD system shows structured activity from an early age (*28*), our observations raise the possibility of a neuronal population that is rigidly wired (*29*) and maintains its organization throughout its lifespan.

Importantly, our data showed how the HD system acquires an alignment at the first visit to the environment and maintains it afterwards. This alignment seemed random and independent of the inputs, as no correlation was found between alignment across animals and environments. Animals were systematically disoriented before each visit. Thus, an association between the sensory inputs and the HD system upon entering the environment may have set an initial alignment, which would have been memorized. Hebbian learning from sensory inputs to a rigidly organized ring attractor neural networks would be sufficient to stabilize an absolute alignment, as suggested in computational models (*17*, *30*) and demonstrated in the fly head-direction system (*31*). Another possibility is that an absolute alignment depends on pre-configured connections from the sensory inputs to the HD system, setting an alignment for a given state of inputs. Such mechanisms will predict that an absolute alignment is maintained across visits, irrespective of the duration between two consecutive visits. However, previous reports have suggested that the alignment is learned (*32*, *33*) Furthermore, the slow drift we observed over weeks rules out the possibility of a rigidly configured system.

Last, the absolute alignment between consecutive visits separated by more than a week did not abruptly change but slowly drifted from their original alignment, increasing with the delay between the two successive visits. This finding suggests that the physiological changes that take place over time, for example, synaptic weight changes between the visual and HD system, do not silently build up to lead to an abrupt remapping of the alignment but instead slowly shift the alignment of the HD system. Note that representation in the visual system also drifts at a faster timescale (*3*), raising the possibility that the HD system must constantly adapt to a changing input.

The HD system is a crucial component of the brain’s navigation system (*34–36*). The PoSub is the main cortical relay of the HD signal and projects directly to the entorhinal cortex (*34*, *35*, *37*), the primary cortical input and output of the hippocampus. The mapping of a familiar environment by hippocampal place cells drifts across sessions (*1*, *8–10*, *13*). Interestingly, our results suggest that this drift occurs irrespective of a stable HD signal. This indicates that, although individual place cells change their firing rate, the global alignment of the hippocampal map should be maintained for several weeks, as suggested by the stability of the position, but not the rate, of each hippocampal place field (*38*). Furthermore, the stability of the HD system begs the question whether spatial representation in the medial entorhinal cortex, especially in the form of a highly organized grid cell system (*39*, *40*), maintain a stable mapping to the environment. Last, these results echo recent computational evidence showing that a cognitive map can emerge in a neural network receiving a stable HD signal, which does not need to be learned end-to-end with the map (*41*).

Experience was essential for maintaining a memory for orientation. Similar observations were made in the olfactory(*2*) and visual systems (*4*). Interestingly, the opposite was observed for hippocampal place cells as experience leads to increased representational drift (*8*, *9*). It is thus possible that experience in a familiar setting, where animals are frequently re-exposed to the same sensory inputs, stabilizes primary sensory representations while increasing the changes in the representation of context by hippocampal neurons.

In conclusion, our findings demonstrate that the HD system is rigidly organized, potentially for the entire lifespan, and forms memories of orientation that last several weeks without re-exposure.

## MATERIALS AND METHODS

### Animals

All experiments were approved by the Animal Care Committee of the Montreal Neurological Institute. Five adult male C57BL/6J mice were used in all experiments. Animals were given water *ad libidum* and kept under a 12h light/dark cycle. Animals were weighed and given food pellets following daily recordings. All mice were maintained at 100 ± 10% of their weight on recording day 1 for the entire recording protocol.

### Viral vectors

Animals were injected with the viral vector AAV2/9-SYN-GCaMP6f (Neurophotonics; batch number 1357). The original concentration was 2.3 × 10^13^ particles ml^-1^. The virus was subdivided into aliquots, stored at −80 °C until use, and diluted in a 1:1 ratio with sterile saline before injection.

### Stereotaxic injections

Animals were anesthetized with isoflurane (5% at induction; 1-2% at maintenance; air-flow 2 L min^-1^) and placed in the stereotaxic frame (Kopf Instruments). Animals were immediately injected subcutaneously with carprofen (20 mg kg^-1^ body weight) and kept on a heating pad for the entire procedure. Eyes were covered with lubricant eye gel (Systane). Fur was removed using Veet hair removal cream and disinfected with 2% Chlorhexidine before applying 5% lidocaine cream to the skin. Surgical instruments were sterilized using a Germinator 500 before opening the skin with a surgical blade. After exposing the skull, a small hole was drilled into the skull over the target site. AAV2/9-SYN-GCaMP6f (100-120 nl) was injected at a speed of 50 nl min^-1^ into the target site using a Nanofil syringe (WPI) and a Pump 11 Elite Nanomite Programmable Syringe Pump (Harvard Apparatus) mounted on the stereotaxic frame. Following injection, the needle was left in place for 10 minutes to allow for proper diffusion and absorption of the virus before slowly removing it from the brain. All animals were injected and recorded from the left hemispheres. Coordinates for injection were AP (from lambda): +0.28 to +0.32 mm, ML (from superior sagittal sinus): −1.19 to −1.15 mm, DV (from dura): −2.20 to −2.45 mm. Furthermore, the needle was titled by 30° in the coronal plane from lateral towards medial.

Following the viral injection and suturing, mice were injected subcutaneously with sterile saline and placed back in their home cage over a heating pad until normal behaviour resumed (max 30 minutes). Animals were subcutaneously injected with carprofen every 24 hours for 72 hours following the procedure and were allowed a recovery period of at least 14 days before further experimentation. Note that animal 2, unlike other animals, was injected and implanted in the same surgery.

### Stereotaxic GRIN lens implantation

GCaMP-injected animals were anesthetized with isoflurane, placed on the stereotaxic frame, injected with carprofen, eyes covered, hair shaved, and skin prepared as described above. The surface of the skull was lightly scratched with a scalpel blade and thoroughly dried before applying and curing three layers of Optibond (Kerr) using UV blue light. A hole was drilled into the skull over the target site, with dimensions large enough to accommodate the 1.0 mm diameter GRIN lens (Inscopix). Following the removal of the meninges, brain tissue above the target region was aspirated using an 18 G needle (BD PrecisionGlide) attached to a suction pump (SHIYN) until the white matter was visible. The GRIN lens was slowly lowered just above the target region using the stereotaxic frame and attached to the skull using a combination of the flowable composite Fusion Flo (Prevest DenPro) and dental acrylic cement (Unifast Trad). Lens implantation coordinates were as follows: PoSub – AP (from Lambda) +0.32 mm, ML −2.35 mm, DV (from skull) −1.3 mm.

A thin layer of dental acrylic cement was then applied to the rest of the skull and a small protective wall was built around the lens to approximately the same height. The wall was filled with Kwik-Cast (World Precision Instruments) to protect the implant.

Animals were injected subcutaneously with sterile saline and kept on a heating pad until normal behaviour resumed. All animals were housed alone following the implant. Additional carprofen injections were delivered daily for a minimum of 4 and a maximum of 7 days post-surgery.

The animals were then base-plated. To this end, after 3-5 weeks to allow for tissue inflammation to completely dissipate, the animals were anesthetized over a heating pad, eyes covered with lubricant gel and placed on the stereotaxic frame. The Kwik-Cast applied in the previous surgery was carefully removed using forceps and the top of the lens was cleaned using 70% ethanol. The Miniscope with a baseplate attached was mounted onto the stereotaxic frame and placed above the lens to image the target tissue and evaluate fluorescence. At this point, animals that did not show any background fluorescence were excluded from the experiment. For animals where fluorescence was observed, the baseplate was cemented to the wall that was made in the previous surgery using dental acrylic cement. After allowing the acrylic to fully dry, the Miniscope was removed from the baseplate and replaced with a baseplate cover. Two layers of black nail polish were then applied to the cement and baseplate to block light. The animals were placed in their home cage over a heating pad until normal behaviour resumed and allowed to rest for 2-3 days before habituation.

### Miniscope recordings

All animals were recorded with the UCLA Miniscope V4. Mice were habituated to being handled by the experimenters for 15 mins per day, for three days before starting the recording protocol. The baseplate cover was removed and the Miniscope was placed into the baseplate while the animals were briefly head-restrained by the experimenter. During the first recording, several different planes of focus were attempted before settling on one for the rest of the experiment. The associated electro-wetting lens (EWL) setting was noted and kept as a reference for subsequent recordings. As an additional measure to ensure recordings focused on the same plane, a screenshot was taken of the chosen plane on day 1, which was then used as a reference when starting all subsequent recordings. Animals were re-plugged with the Miniscope or the EWL setting adjusted as necessary to match the reference image.

The animals were recorded in four different environments: Circular, Square, Radial (8-arm) maze and Open field (1m^2^ square arena). The Circular, Square and Radial maze environments were all located in different corners of the same room, separated by black curtains. In contrast, the Open field was located in a separate room, on a different floor of the building. Animals were recorded during freely moving foraging behaviour on weekdays in 20-minute periods. Reflective markers were permanently mounted on the Miniscope. The location and head direction of the animals were tracked with a multi-camera (6 to 8) system (Optitrack) in all four environments.

The video recordings from the Miniscope were concatenated and processed for motion correction using NoRMCorre (*42*). Regions of interest for each recording were then detected using CNMF_E (*43*). ROIs were aligned or registered across days using CellReg (*26*).

### Silicon Probe Implantation

Animals were anesthetized using isoflurane and placed on the stereotaxic frame as previously described. A silver wire was inserted into the cerebellum to serve as a reference. Two types of silicon probes were used for these experiments, one containing a single shank (Cambridge NeuroTech H5) and one containing two shanks (Diagnostic Biochips 128-3). The probes were mounted on a moveable microdrive. A hole was drilled into the skull above the target site large enough to accommodate the size of the probe, about 0.1 mm on each side of the craniotomy. The probe was slowly lowered above the target site and secured to the skull using a light-cure adhesive (Kerr OptiBond) and dental acrylic cement (Unifast Trad). Implants included either tetrodes or silver wires directed at CA1. A copper mesh was attached with a flowable composite around the probe to reduce electrical noise in the recordings. Animals were injected subcutaneously with saline (around 0.8 ml) and carprofen (20 mg kg^-1^) and placed on a heating pad until normal behaviour resumed. Animals were housed individually following the implant surgery. Carprofen injections were delivered daily for at least three days post-surgery. Silicon probe implantation coordinates were as follows: Angled PoSub implant (at a 30° angle to the right) – AP (from Bregma) −3.70, ML −1.10, DV −1.50. Straight PoSub implant – AP (from Lambda) +0.30, ML - 2.35, DV – 1.10. CA1 implants – AP (from Bregma) −1.80, ML −1.20, DV −1.10.

### Electrophysiological Recordings

Animals were monitored daily and allowed to recover for 5-7 days post-surgery prior to recording. The probe was slowly lowered into the target region in steps of 0.03 mm over a few (3–4) hours. Animals were recorded for 15-20 minutes during free foraging behaviour in 3 different environments: Circular, Square and Radial maze. The mice were placed in their home cage, covered, and disoriented for 2-5 minutes in between environments to prevent associations between them. The neurophysiological signals were acquired at 20 kHz using the Intan RHD 2000 Recording System. The signal was then processed using an automated spike sorting algorithm (Kilosort2.5). Isolated units were then manually curated using the Klusters software. The animals’ location and head-direction were tracked using the same multi-camera system and reflective markers as used in the calcium imaging experiments.

### Inclusion and exclusion criteria

To track the activity of the same units over time, the field of view must remain the same, meaning that the same Miniscope must be used. For animal #2, the original Miniscope used to record in the circular environment broke, leading to a shift of the field of view that rendered us unable to track most of the ROIs in the original plane. For this reason, this animal was excluded from the analysis of between-rig comparison of angular offset (Fig. 2). Given some ROIs were retained from the original field of view, the animal was still included in the analysis of between-rig remapping (Fig. 2) and memory stability (Fig. 3). The field of view was lost between multi-environment recordings and probe trials for animal #5, leading to the exclusion of that animal for the analysis of environment-specific directional stability (Fig. 3). Only animals #1, #3, #4 were recorded for the comparison of one and four-week long directional stability (Fig. 4).

**Figure 4:**
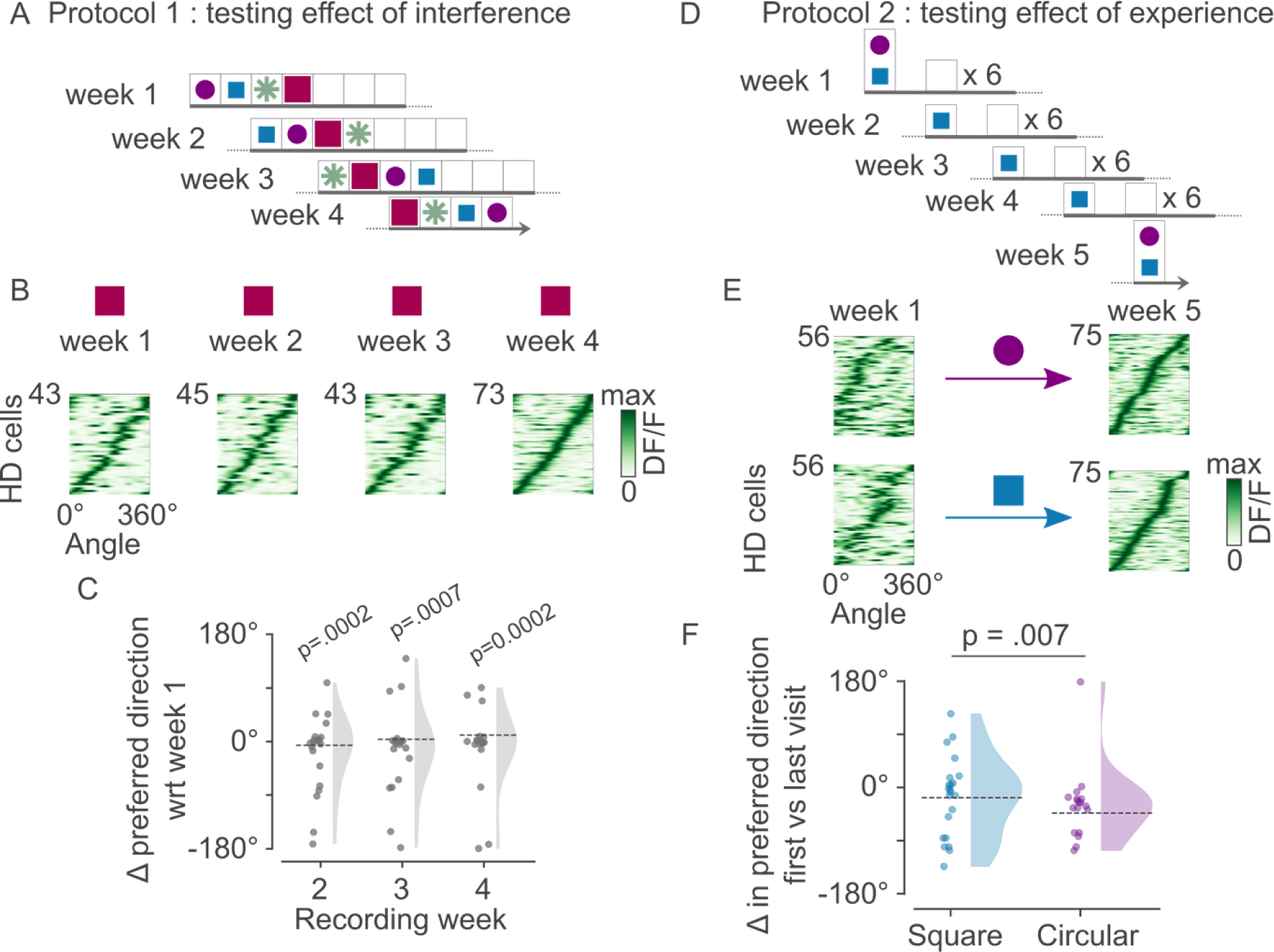
Lack of experience, but not interference, explains slow drifts in memory of orientation. A) Hypothesis 1: Experience of other environments interferes with orientation memory. For four weeks, familiar environments are visited every week in a different order (16 sessions total). B) Tuning curves of a population of HD cells, for an example animal in the open field environment across the four weeks, sorted by week 4. C) Change in preferred direction with respect to the first visit of each environment. All environments and all animals pooled (V-test, not different from 0°; p<0.001 for all weeks; n = 20, 20, 18 sessions per week respectively; 5 animals). D) Hypothesis 2: Delay between successive experience accounts for drift in orientation memory. Two familiar environments were visited in a single day, followed by weekly visits of one environment for three weeks, after which both environments were revisited in a single day (7 sessions, 5 weeks). E) HD cell population tuning curves for the first versus last week in an example animal, sorted by week 5 for both environments. F) Change in preferred direction between the first and last exploration in the same environment (p = .007, Kolmogorov-Smirnov two-sample test; n = 34 HD cells; 3 animals).

**Table 1:** Summary of dataset.

| Animal # | Experimental # | Age at time of first recording | Age at time of last recording | Total number of sessions recorded | Average ROIs recorded ± 1 STD |
| --- | --- | --- | --- | --- | --- |
| 1 | A0667 | 19 weeks | 43 weeks | 82 | $65 \pm 19$ |
| 2 | A0662 | 29 weeks | 44 weeks | 72 | $51 \pm 11$ |
| 3 | A0668 | 19 weeks | 43 weeks | 82 | $186 \pm 38$ |
| 4 | A0670 | 19 weeks | 43 weeks | 82 | $60 \pm 10$ |
| 5 | A0642 | 34 weeks | 61 weeks | 70 | $119 \pm 30$ |

### Data analysis

All analyses were done using customized code written in python (3.8) with the following libraries: pandas, numpy, scipy, scikit-learn, pycircstat and pynapple (*44*).

### Head-direction cell classification - Miniscope

For each session, the animal’s head direction in the horizontal plane was extracted from the multi-camera tracking system (Optitrack). HD tuning curves (i.e. average DF/F) were computed in 3° bins, divided by the total time spent by the animal in each angular direction. The resulting tuning curves were then smoothed using a Gaussian kernel (σ = 12° bins). The correlation coefficient was computed between tuning curves generated from the first versus the second half of each session. ROIs with correlation values below 0.4 between the two halves were discarded. Head-direction information (HDI) was then computed for each ROI and each session (*45*). ROIs with an average HDI above 0.1 (across sessions) were labelled as HD cells. While all HD cells were shown in example population tuning curves, sessions where the cell was detected but the HDI fell below 0.11 were discarded from further analysis. Furthermore, only HD cells that were tracked on multiple days and with cell registration confidence scores above 0.8 (from the *CellReg* software) were included in further analyses. Given the long timescale of Fig. 3, a score above 0.6 in these analyses was acceptable. ROIs had to be detected for at least six sessions for the analyses in Fig. 1, 2 and 4A-C. As the total number of sessions was smaller for analyses in Fig. 3 and 4D-F, and Supplementary Fig. 6E,F, the number of sessions in which ROIs had to be detected was set to 2.

### Head-direction cell classification – Electrophysiology

Neurons with firing rates above 1 Hz and an HDI above 0.3 were classified as HD cells. Note that HDI values cannot be directly compared between the imaging and electrophysiology, explaining the different thresholds between the two datasets.

### Dimensionality reduction and population data visualization

To visualize the organization of neuronal activity at the population level, we projected the data on a two-dimensional subspace using ISOMAP projections (*46*) (using the *scikit-learn* library). HD neurons were selected according to the correlation value of their tuning curves (>0.8) and HDI (>0.1) in individual sessions, as previously described. Unlike tuning curves for which continuous DF/F was used, here we focused the analysis on calcium transients to reduce the effect of residual correlation resulting from the low-frequency dynamics of calcium fluorescence. Specifically, DF/F transients were smoothed with a Gaussian kernel (σ = 133 millisecond bins) and then normalized. The transients were then downsampled by taking the average fluorescence in 0.25-second bins. Timepoints when the average of the normalized binned transients fell below 0.1 were discarded. The remaining binned transients were projected to a two-dimensional plane using ISOMAP (*46*). The number of neighbors was set to 50.

For the electrophysiological recordings, spike rate was computed in bins of 0.2 seconds and smoothed with a Gaussian kernel (σ = 0.8 second bins). Only epochs during which the animal’s linear velocity was above 2 cm second^-1^. The remaining bins were projected using ISOMAP as above for calcium transients.

### Jaccard Score

To evaluate the level of registration across days, we used the Jaccard Score, which measures the similarity between two sets of pairs of binary vectors corresponding to the presence or absence of HD cells on a given day. A value of 0 indicates no overlap/similarity between the binary vectors, while a value of 1 indicates the two vectors are duplicates of one another. This analysis was restricted to HD cells as described above. For Fig. 1G and Supplementary Fig. 2D and 3A,B; the median Jaccard index was calculated for all pairs of recordings that matched the relevant day difference.

### Change in the preferred tuning of HD cells over days

For the comparison of a pair of recordings, only HD cells detected in both recordings were included. The tuning curves of the same HD cell were cross-correlated. The angular shift corresponding to the maximum correlation value was taken as the angular difference in preferred directions for that HD cell across the environment. The circular mean of the angular difference of all HD cells was computed as thpee change in preferred tuning for each session.

### Change in the angular offset of HD cells across days

For each pair of HD cells, only recordings where both HD cells were detected were included. The tuning curves of each pair of HD cells were cross-correlated with each other to determine the angular offset. The change in angular offset was then computed as the minimum circular difference between angular offset values across pairs of days. Corresponding shuffles were computed by shuffling HD cell pair identity within a single day prior to computing the change in angular offset, 1000 times.

### Orientation remapping and mean vector length

Orientation remapping was calculated in a similar way as the change of preferred HD tuning over days. The remapping for a pair of environments was computed as the circular mean of the angular shift of all HD cells detected in both recordings. The average remapping from one environment to the next was calculated as the circular mean of the average remapping of all pairs of recordings between the two environments. To determine the coherency of remapping, the mean vector length was computed using each pair of recordings and compared to shuffles.

To compare the probe trial and the previous 2-week-long visit (Fig. 3), the remapping was computed using only the last visit of each environment against the probe trial visit.

Corresponding controls were computed by shuffling HD cell identity within a single day prior to computing the change in preferred direction between days, 1000 times.

## ACKNOWLEDGMENTS

We would like to thank Lynda Mainville for technical support. We are thankful to Mark Brandon, Daniel Levenstein and other members of the Peyrache laboratory for comments on the earlier version of the manuscript.

This work was supported by the Canadian Research Chair in Systems Neuroscience (AP), CIHR Project Grant 155957 and 180330 (AP), NSERC Discovery Grant RGPIN-2018-04600 (AP), Canada-Israel Health Research Initiative, jointly funded by the Canadian Institutes of Health Research, the Israel Science Foundation, the International Development Research Centre, Canada and the Azrieli Foundation 108877-001 (AP), the New Frontiers in Research Fund Exploration grant NFRFE-2021-00926 (AP), HBHL - Healthy Brains for Healthy Lives Innovative Ideas grant 1c-II-15 (AP), and Vanier CGS (SSC).

## AUTHOR CONTRIBUTIONS

Conceptualization: SSC, GV, AP; Methodology: SSC, GV, AP; Software: SSC, GV, AP; Validation: SSC, GV, AP; Formal analysis: SSC, GV, AP; Investigation: SSC, GV, AP; Resources: AP; Data Curation: SSC, GV; Writing - Original Draft: SSC, AP; Writing - Review & Editing: SSC, AP; Visualization: SSC, AP; Supervision: AP; Project administration: AP; Funding acquisition: AP.

## Supplementary Information

**Supplementary Fig. 1:**
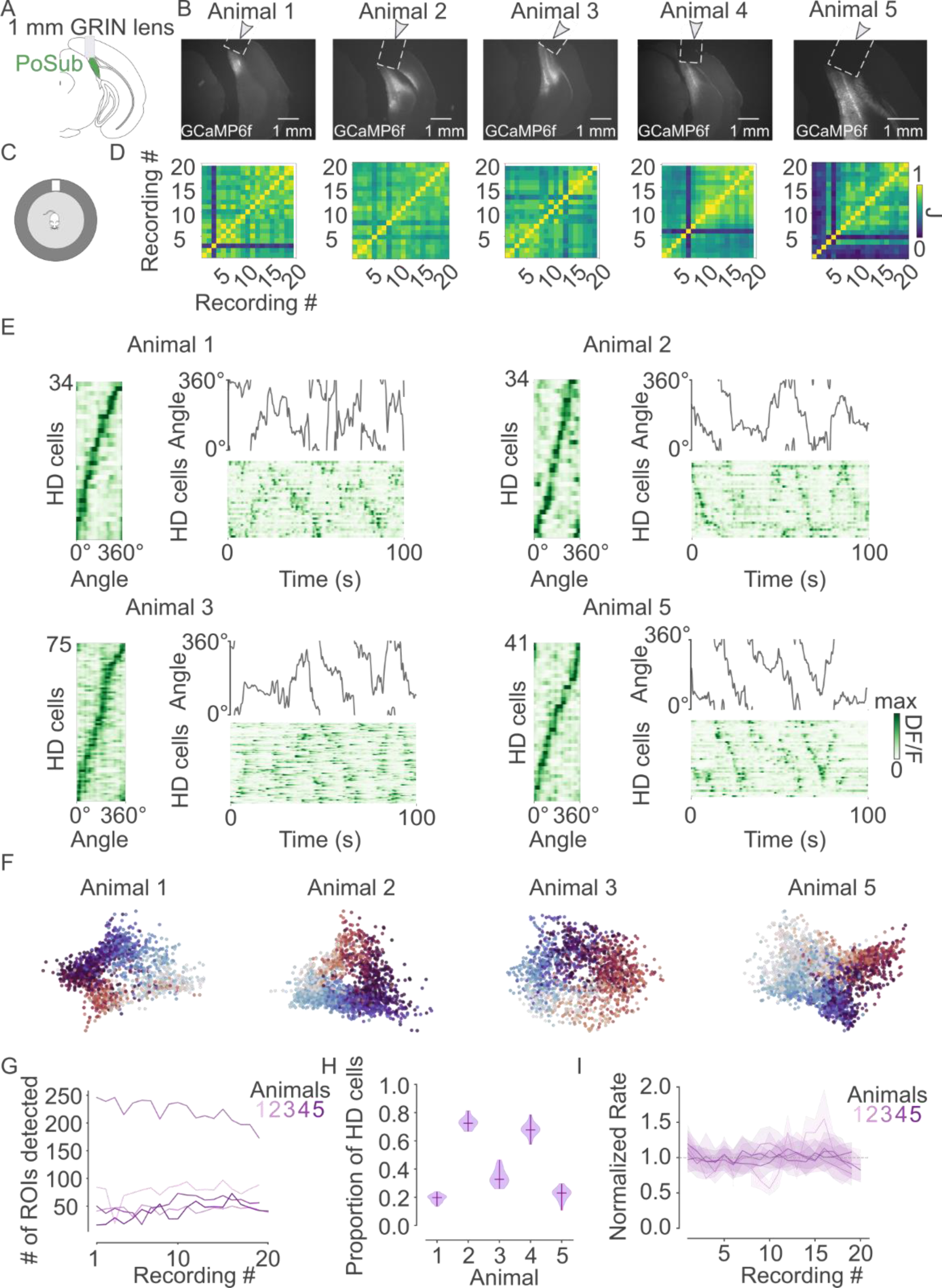
Longitudinal recordings in the PoSub. A) GRIN lens position above PoSub. B) Example histology of each animal. C) Schematics of the arena in which freely moving recordings were performed. A white card was placed on the wall and was the main visual cue visible to the animal. D) Day-to-day Jaccard index for all animals, showing the similarity of HD cell registration across sessions. E) Color-coded tuning curves of all HD cells simultaneously detected as well as calcium transients from an example recording session, for all animals (except animal #4 shown in Fig. 1). F) Low-dimensionality projection of HD cell population calcium transients from an example session from each animal (except animal #4 shown in Fig. 1) (see Methods). G) ROI count per animal across all recording sessions. H) Proportion of ROIs detected as HD cells throughout the recording protocol. I) Rate of HD cell calcium transients (detected as peaks in calcium fluorescence) for all animals across recording sessions, normalized by the mean across sessions.

**Supplementary Fig. 2:**
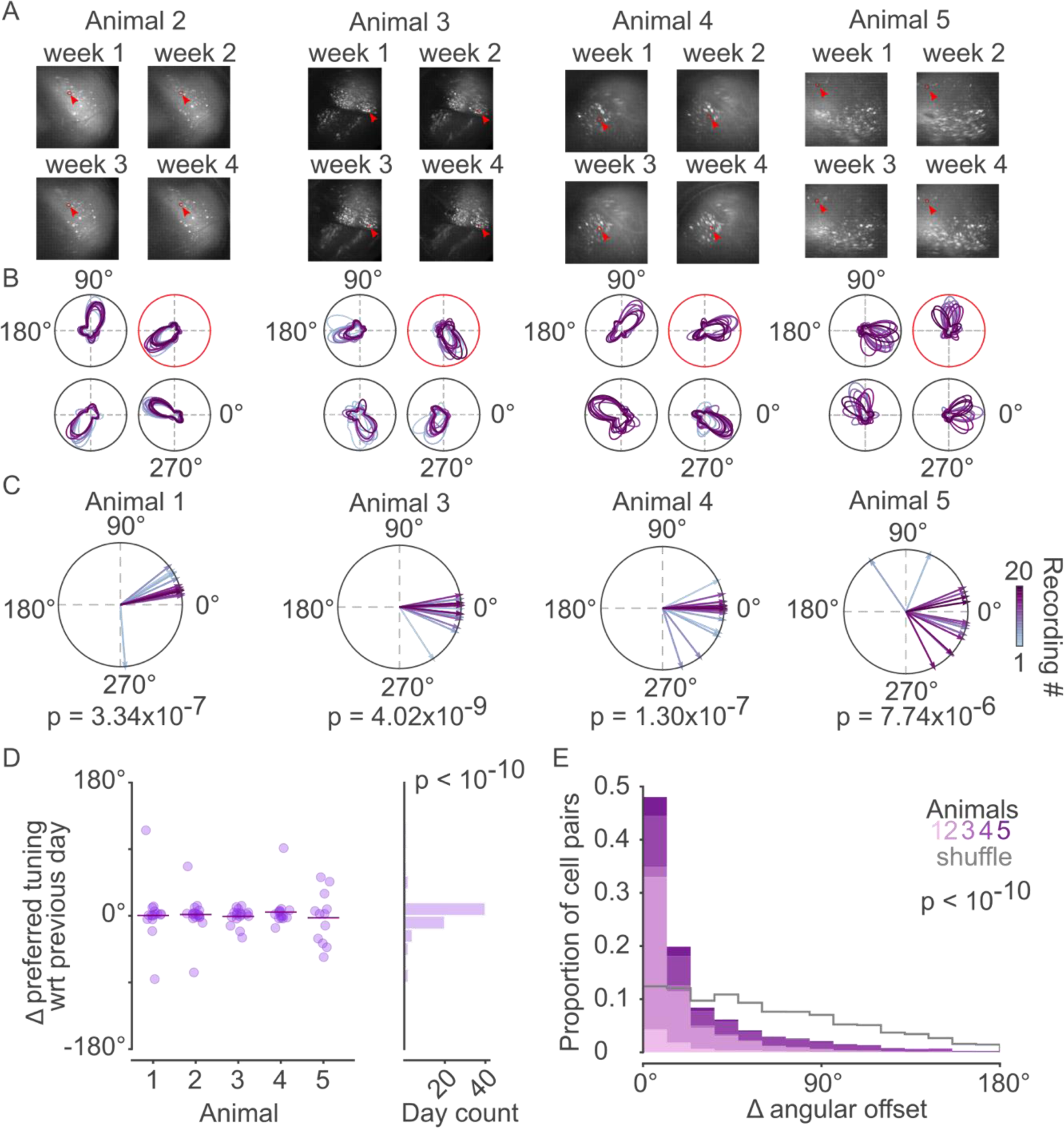
Long-term stability of HD cell direction tuning. A) Spatial footprint of the circled ROI in panel B for each animal not in Fig. 1, in four sessions across the four weeks of the recording protocol. B) Tuning curves of four example ROIs from each animal not in Fig. 1 across all sessions where the ROI was detected. C) Population average of across-session angular difference in preferred direction, relative to the last session for all animals not in Fig. 1 (V-test, not different from 0°, p<0.001 for all animals, n = 16, 18, 16, 16 sessions respectively). D) Distribution of population-wise angular difference relative to the previous session for all five animals (V-test, not different from 0°, p<10^-10^, n=75 sessions) E) Difference in angular offset across all HD cell pairs and all recording pairs in the recording protocol, shown for individual animals relative to control datasets of shuffled cell pair identities (p < 10^-10^, Kolmogorov-Smirnov two-sample test on pooled animals, n= 25,901 pairs)

**Supplementary Fig. 3:**
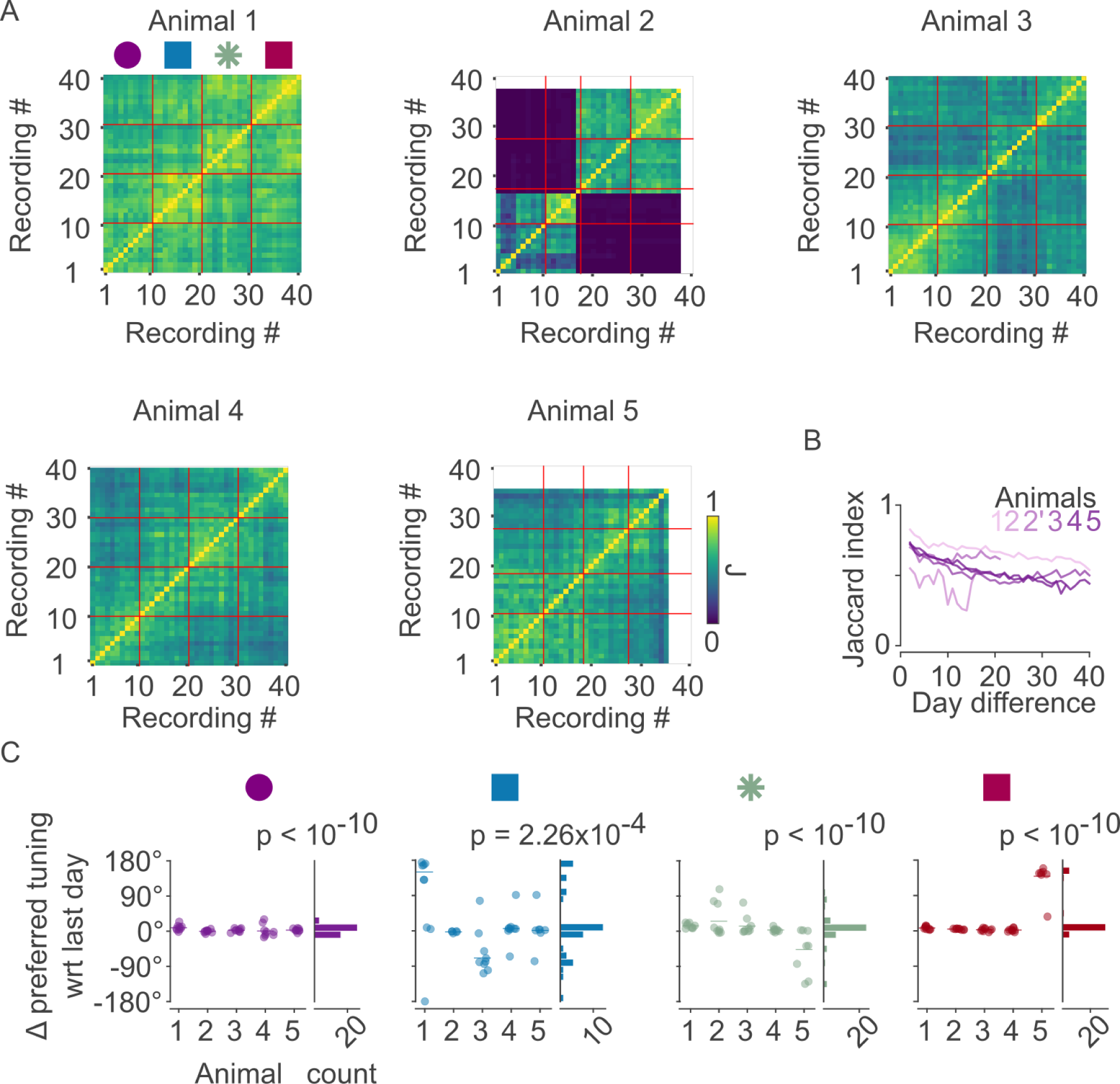
Longitudinal PoSub recordings in multiple environments. A) Day-to-day Jaccard index for all animals, showing the similarity of HD cell registration across sessions. Red lines indicate the change from one environment to the next. Note that miniscope was changed between 2^nd^ and 3^rd^ environment for animal #2 (see Methods). B) Average Jaccard index as a function of duration between pairs of sessions for all animals. Data for animal 2 are split between the two epochs for which different miniscopes were used. C) Stability within each environment with respect to the last day (V-test, not different from 0°, p < .001 for all environments, n = 45, 39, 44, 43 sessions respectively). To note, for animal 2 in the square environment the next-to-last day was used due to poor cell registration on the last day. Repeated-measures ANOVA across environments reveals no significant difference between environments. Furthermore, note that the apparent instability in the square environment mostly resulted from 90° and 180° remapping, certainly as a consequence of the environment geometry.

**Supplementary Fig. 4:**
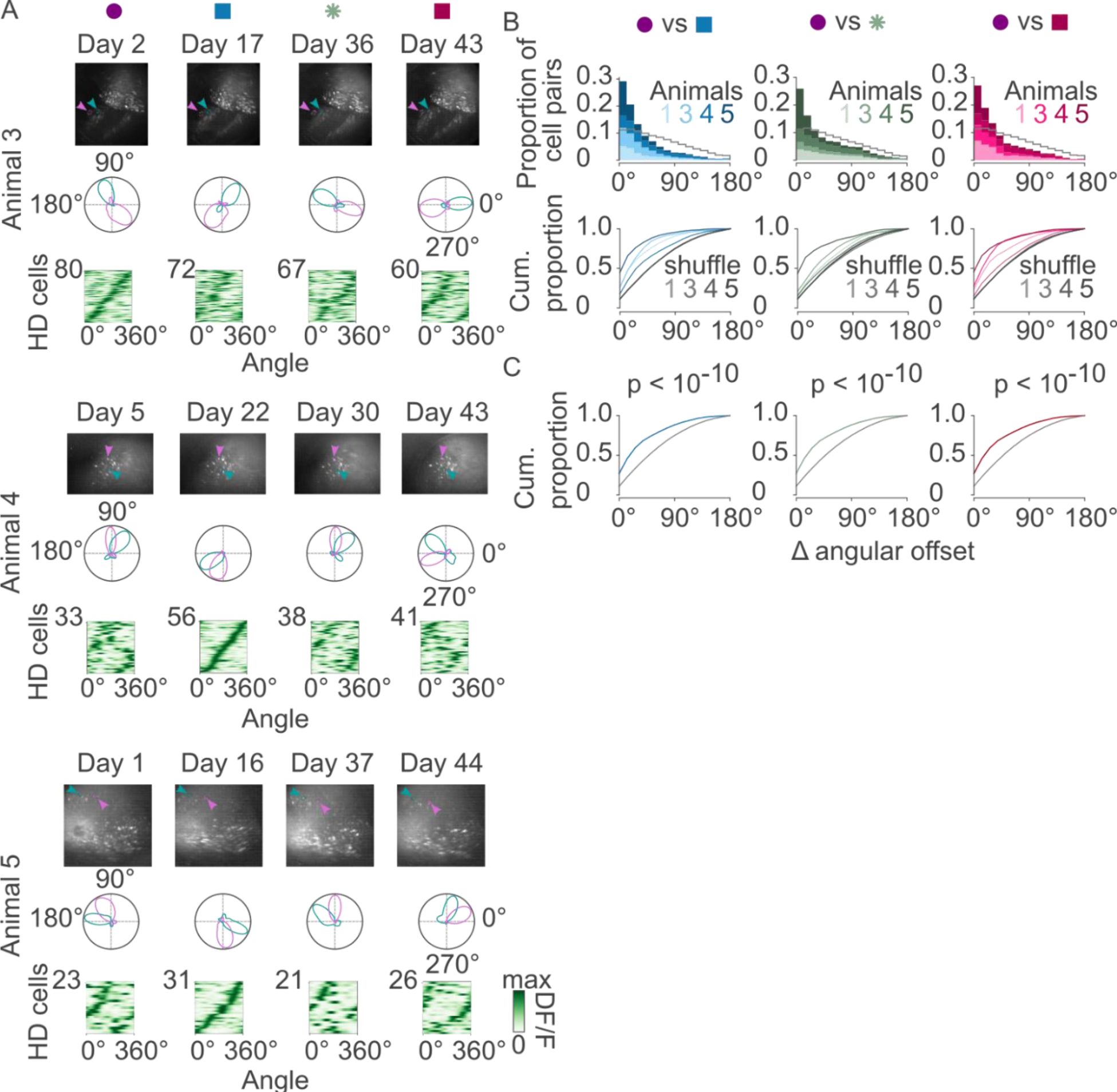
Long-term and across-environment stability of the HD cell population. A) Additional examples of month-long stability except for the animal in Fig. 2. For each animal, one day from each environment is shown. Top, example field of views across the recording protocol with two identifiable HD cells in each animal. Middle, example tuning curves of the two HD cells shown in the field of view. Bottom, tuning curves of all HD cells detected in these example days, sorted with respect to the day with the highest number of HD cells. B) Difference in angular offset across all HD cell pairs, for the first week of exploration of the circular environment versus the two weeks of exploration of the three other environments. Top, angular offset differences for individual animals; bottom, cumulative proportion shown for each animal and their respective shuffle. C) Same as panel B, shown for the pooled population relative to control datasets of shuffled cell pair identities (p < 10^-10^, Kolmogorov-Smirnov two-sample test, n = 9,430; 6,400; and 6,160 pairs respectively).

**Supplementary Fig. 5:**
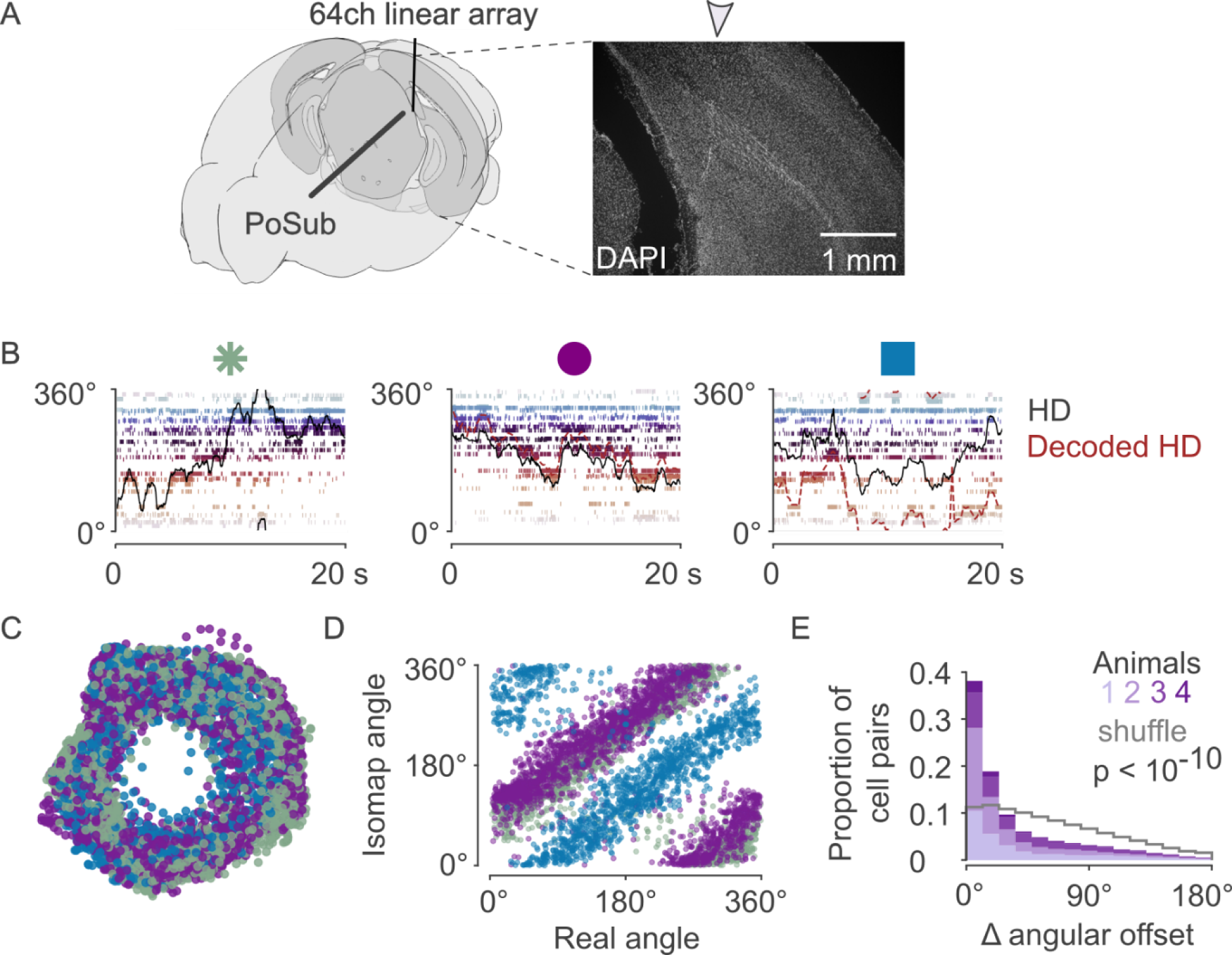
Electrophysiological recordings in the PoSub across environments. A) Diagram of linear electrode array into the PoSub (left) and example histology (right) illustrated with brainrender (*27*) B) Raster plots from one example animal exploring three environments. Neurons are ordered and colored according to preferred HD in the 8-arm maze. Bayesian decoder (*47*), trained on the exploration in the 8-arm maze. C) Low-dimensional projection of HD cell population activity across all three environments (same color code as environments shown in panel B). Each point corresponds to one time bin. D) Isomap-defined angles (i.e. angular value in polar coordinates for each point in panel C) as a function of actual animal’s head-direction across the three environments. E) Difference in angular offset across all HD cell pairs and across all environment comparisons, shown for individual animals relative to control datasets of shuffled cell pair identities (p < 10^-10^, Kolmogorov-Smirnov two-sample test, n = 18,450 pairs).

**Supplementary Fig. 6:**
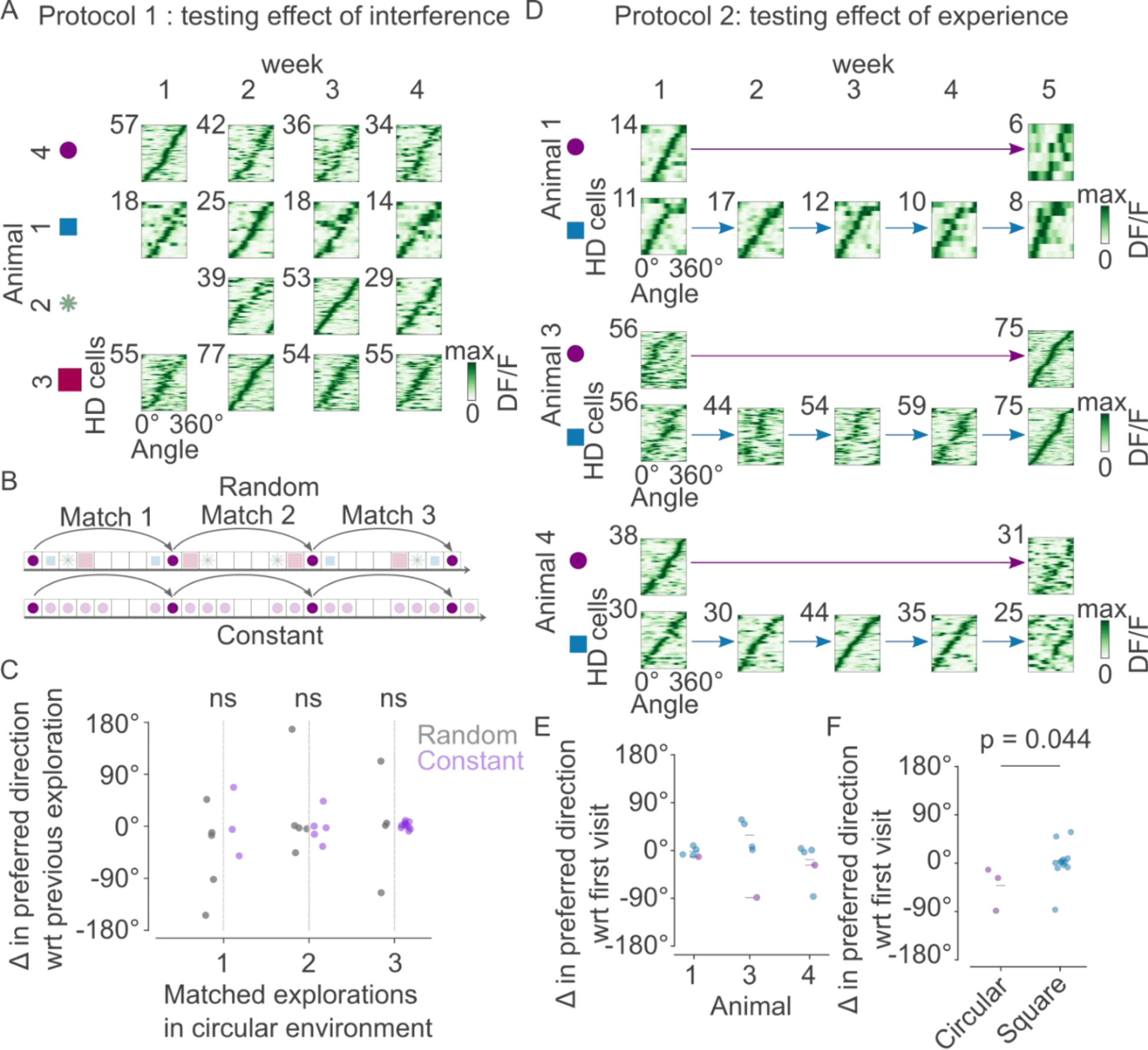
Orientation memory stability. A) Additional examples of animals and environments across the four weeks of recordings (similar to Fig. 4B), sorted with respect to the week with the most HD cells. B) Matched explorations between the random environment exploration (shown in Fig. 4) and the constant four-week visits to the circular environment (shown in Fig. 1). Sessions were matched according to number of days between exploration and recording weeks in the circular environment. C) Comparison of matched explorations showed no significant difference between the random and constant environment experiments (p > 0.05, Kolmogorov-Smirnov two-sample test; n = 9, 13, 16 sessions respectively). D) All animals and recordings across the five weeks of recordings (similar to Fig. 4E), sorted with respect to the week with the most HD cells for each environment. E) Change in preferred direction with respect to the first visit of each environment, per animal. Sessions are color coded according to the environment color in panel D. Mean change per environment shown by a colored bar. F) Change in preferred direction with respect to the first visit of each environment showed a significant difference between the circular and square environments (p > 0.044, Kolmogorov-Smirnov two-sample test, n = 15 sessions).

